# Mangrove forests mitigate coral bleaching under thermal stress from climate change

**DOI:** 10.1101/2021.06.04.447049

**Authors:** Jack V. Johnson, Jaimie T.A. Dick, Daniel Pincheira-Donoso

## Abstract

Anthropogenic marine heatwaves are progressively degrading coral reef ecosystems worldwide via the process of coral bleaching (the expulsion of photosynthetic endosymbionts which reveals the coral skeleton). Corals from mangrove lagoons are hypothesised to increase resistance and resilience to coral bleaching, highlighting these areas as potential natural refuges for corals. Our study, the first conducted at a global-scale, reveals that coral reefs associated with mangrove forests are less likely to bleach under thermal stress, and thus, under scenarios of climate warming. The onset of severe bleaching occurred after 3.6 Degree Heating Weeks (DHW) in mangrove-associated reefs, compared to 2.23 DHW for non-mangrove associated reefs. These findings highlight the critical role of mangrove forests for coral reef persistence under climate change. Accordingly, conservation actions targeting the protection of mangroves are expected to contribute to the resilience and resistance of reef corals from bleaching as marine heatwaves continue to become more common.

## MAIN TEXT

### Introduction

Coral reefs are continuing to decline globally, primarily driven by anthropogenic heating (*1*–*3*). Increased sea water temperatures and marine heatwaves are inciting coral bleaching, a process whereby photosynthetic endosymbionts are expelled by the cnidarian host (*4*–*8*). Continued coral bleaching is increasingly leading to mortality of coral colonies over entire reefs (*1*, *9*, *10*), critically threatening the key ecosystem services provided by coral reefs which host ∼25% of marine biodiversity and support the livelihoods of over half a billion people (*11*, *12*).

Given the complex interactions between the environmental conditions at local scales with the coral holobiont (i.e. the coral polyp, endosymbionts and microbiota) (*13*), the degrees of bleaching from different coral taxa vary across different regions of the world, creating spatial patterns of bleaching (*14*–*16*). In addition, coral bleaching can often be a sub-lethal process to aid individuals for withstanding environmental stress (*7*). However, given the increased frequency and intensity of marine heatwaves which induces bleaching over mass scales, leading to high levels of mortality(*1*, *9*, *10*), functional and taxonomic homogenisation (i.e. reef flattening) of entire coral reef ecosystems is occurring (*2*, *9*, *10*). Therefore, elucidating patterns of spatial variation in coral susceptibility to bleaching, and the underlying mechanisms responsible, is critical for the identification of factors and ecological settings with the potential to mitigate the decline of coral reefs, which can in turn translate into the design of conservation actions.

In particular, mangrove lagoons have been suggested to act as potential refuges for coral reefs, either by maintaining favourable environmental conditions (*17*), or by preconditioning corals to future environmental stressors (*18*). Mangrove forest lagoons appear to mitigate the severity of coral bleaching under thermal stress given that they are shaded environments, often with higher turbidity levels which reduce solar irradiance and temperature – potentially similar to that of near-shore reefs with reduced levels of bleaching under thermal stress (*19*–*21*). Furthermore, coral holobionts within mangrove lagoons show unique microbial compositions allowing them to survive in extreme environments which may represent future climate conditions (*22*, *23*). Therefore, coral reefs associated with mangrove forests may be more resilient or resistant to the synergistic effects of factors that trigger bleaching in exposed marine areas as they have been preconditioned to environmental stress (*18*, *23*). Concurrently, these corals may proliferate further out onto coral reef environments, displacing corals more susceptible to thermal stress (*24*). As rising global temperatures continue to increase marine heatwaves (*25*) and thus bleaching intensity (*1*), taxonomic displacement of corals with low thermal tolerance by preconditioned coral species from chronic stress becomes more likely (*16*, *24*, *26*).

Despite emerging evidence of mangrove-associated corals showing higher resilience and resistance to thermal stress at local-scales (*18*, *23*), the generality of the impact of mangroves as environments that buffer bleaching remains unknown as global-scale analyses are lacking. In this study, we implement the first global-scale analyses (Fig. 1) aimed to address the hypothesis that coral reefs associated with mangrove forests show resistance to bleaching under thermal stress. To test this hypothesis, we used 9,456 Reef Check Surveys across 3,686 sites from 2002-2017 to determine bleaching responses under thermal stress associated with increased mangrove area.

**Fig 1.**
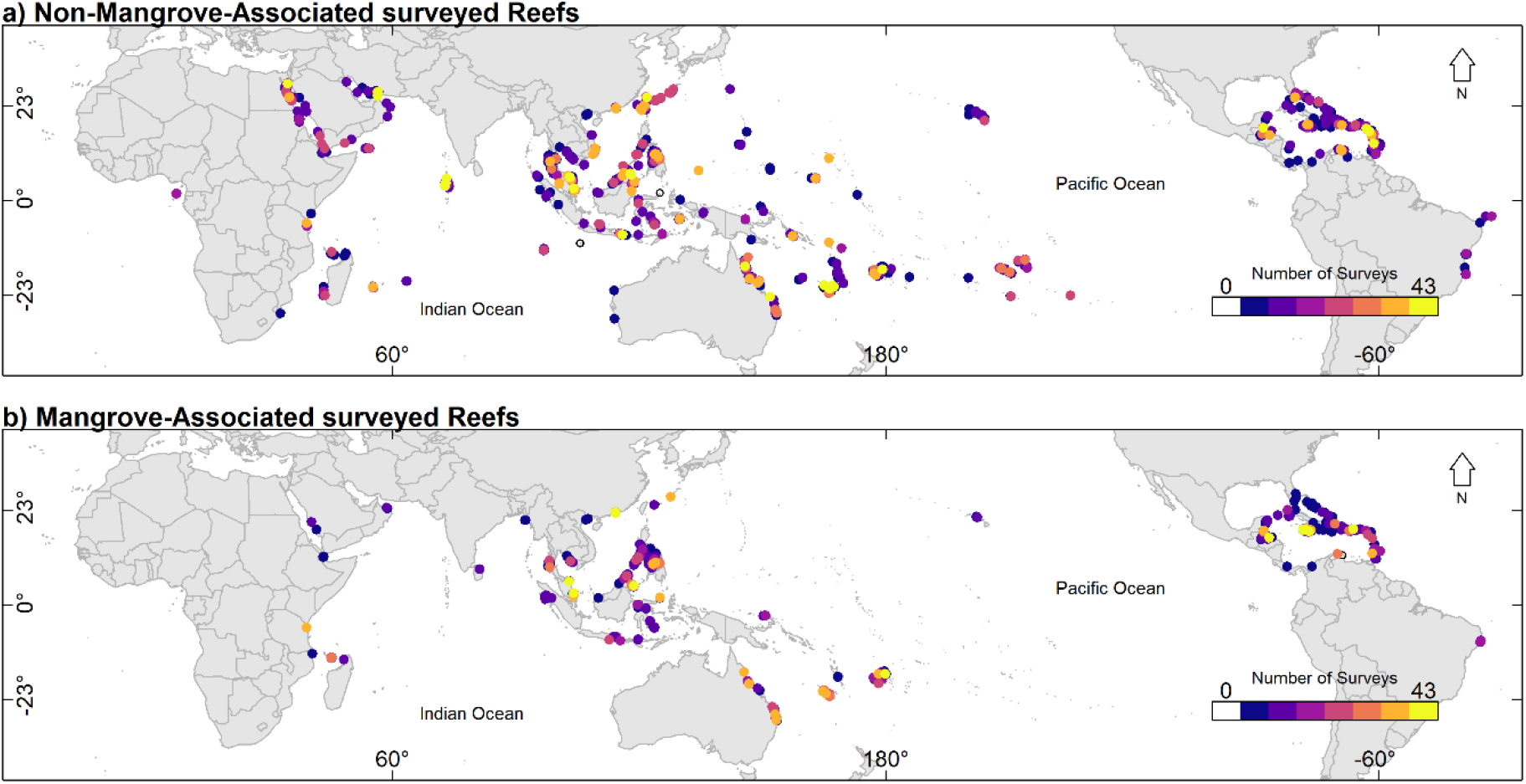
Reef Check Surveys. Global distribution richness of the number of reef check surveys undertaken at each site used in this study. a) Are reef check surveys which are not within a 4km range of mangrove forests while b) are sites surveyed within a 4km radius of mangrove forests.

We used a distance of 4km from each reef check survey to determine the mangrove area associated with each reef check survey, as determined by sensitivity analysis (Table 1), and assessed the interaction between the cumulative numbers of Degree Heating Weeks (DHW) with mangrove area at each Reef Check survey site.

**Table 1.** Sensitivity analysis. Mean model coefficients with upper and lower Credible Interval (CI) coefficients, and deviance information criteria (DIC) scores from the Bayesian ordinal regression models ran as sensitivity analysis.

| Distance | variable | Mean | Lower CI | Upper CI | DIC |
| --- | --- | --- | --- | --- | --- |
| 1km | DHW | 0.322307 | 0.286019 | 0.357965 | 15054.89 |
|  | Mangrove area | 0.057515 | -0.00217 | 0.115447 |  |
|  | Interaction | -0.02133 | -0.06539 | 0.024857 |  |
| 2km | DHW | 0.331814 | 0.290522 | 0.371336 | 15126.88 |
|  | Mangrove area | 0.169023 | 0.113326 | 0.223857 |  |
|  | Interaction | -0.0399 | -0.08433 | 0.004447 |  |
| 4km | DHW | 0.358085 | 0.316068 | 0.399579 | 15015.18 |
|  | Mangrove area | 0.173987 | 0.117529 | 0.231106 |  |
|  | Interaction | -0.07611 | -0.12162 | -0.03147 |  |

## Results

### Bleaching responses

The Bayesian hierarchical ordinal regression model shows that the interaction between DHW and mangrove area reduce the probability of coral bleaching globally (Fig. 2).

**Fig 2.**
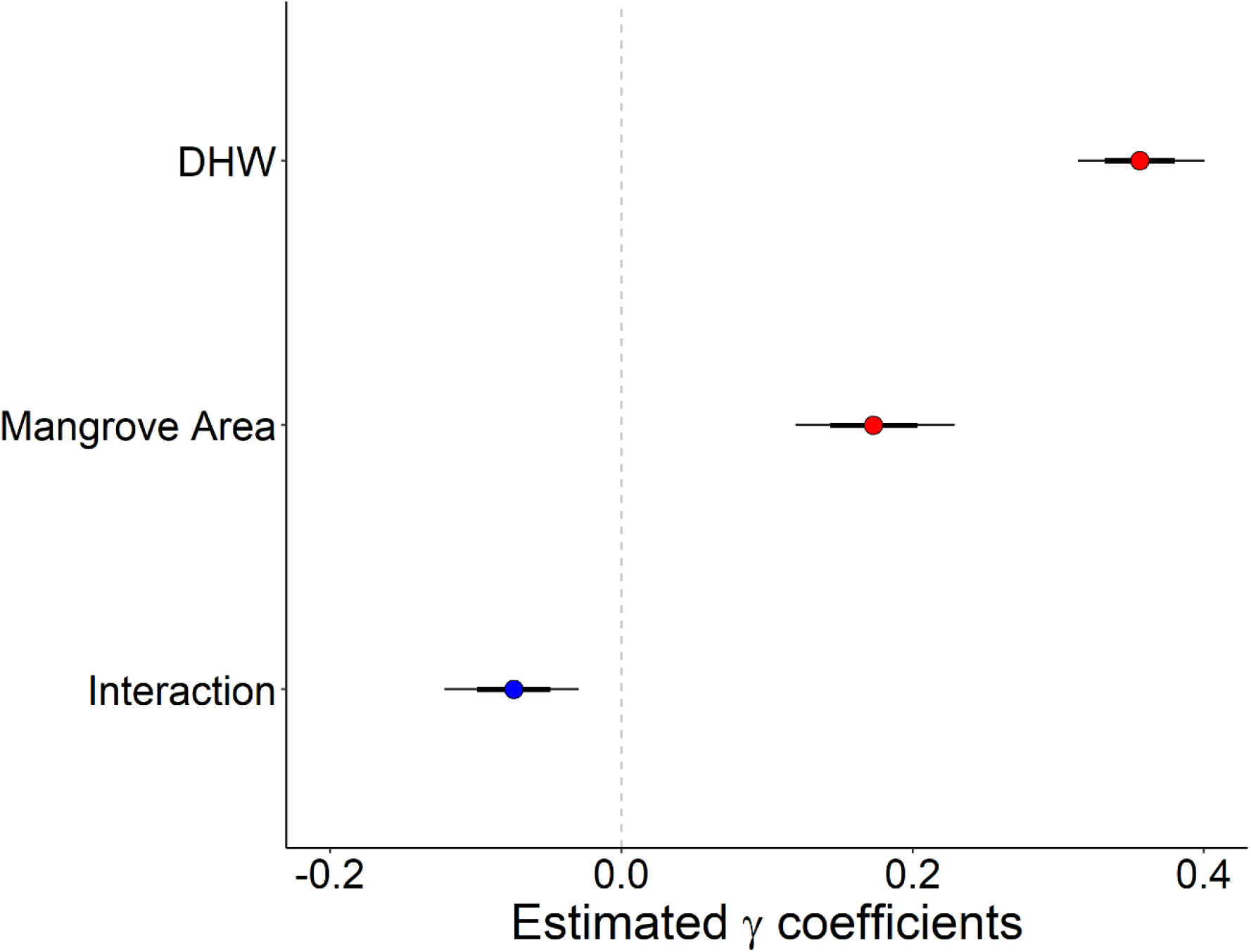
Model outputs. Bayesian hierarchical ordinal regression model showing the relationship between bleaching severity with the mangrove area within a 4km radius of surveyed reef, degree heating weeks (DHW), and the interaction between mangrove area and DHW. The circle represents mean values, thick horizontal line represents 50% credible intervals, thin horizontal line represents 89% credible intervals. Results are considered significant if the 89% credible interval line does not cross zero (54). Positive gamma coefficients show an increase in bleaching likelihood (red dot) while the negative coefficient (blue dot) represents a decrease in bleaching likelihood. Data were from 9,456 reef check surveys from 2002-2017 covering 3,686 sites.

Meanwhile, increasing DHW alone increased the probability of coral bleaching, as did an increase in mangrove area alone. Therefore, under elevated thermal stress, coral bleaching is less likely at coral reef sites associated with mangrove forests.

### Bleaching thresholds

When comparing the raw DHW values where the onset of bleaching occurred, our analyses revealed no significant difference between mangrove and non-mangrove associated sites. Furthermore, the onset of bleaching for the mild and moderate severity bleaching categories did not occur at significantly different DHW thresholds between mangrove and non-mangrove sites. However, the onset of severe bleaching occurred after 3.6 DHWs at mangrove sites, which was significantly higher (Pr(>X^2^) < 0.05) than the 2.23 DHWs required to induce severe bleaching at non-mangrove sites (Fig. 3). Therefore, mangrove sites are more resistant to severe bleaching under elevated thermal stress from climate change.

**Fig 3.**
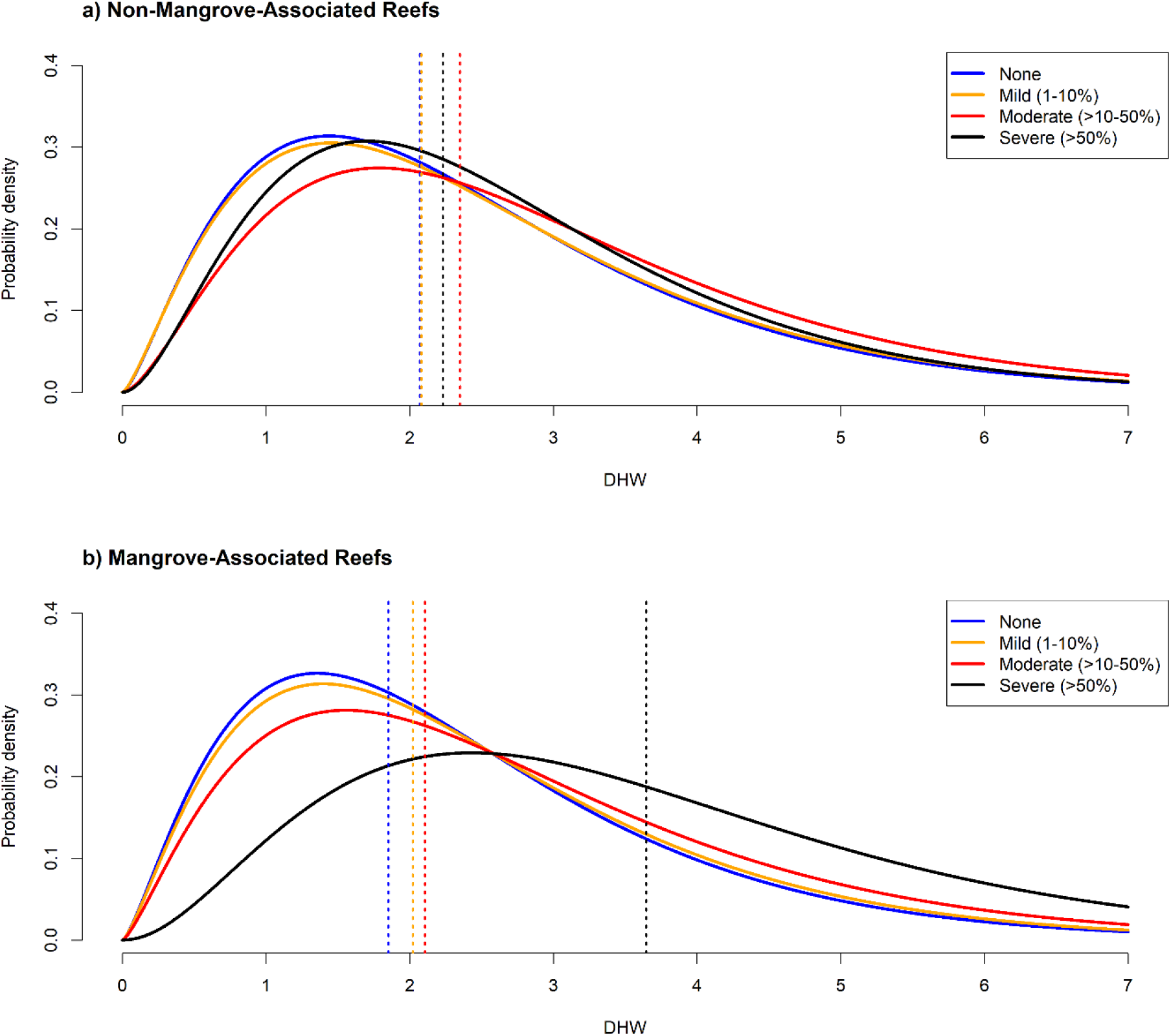
Bleaching thresholds. Probability densities of the degree heating weeks (DHW) where the onset of bleaching occurs for each severity category. a) Represents non-mangrove associated reefs, b) represents mangrove associated reefs. Solid curved lines represent gamma best fit probability distributions with dashed vertical lines representing the median number of DHWs where the onset of each bleaching category occurs. Gamma model outputs for a) and b) are in Table S3.

## Discussion

Our study provides the first global-scale analysis of the role that mangroves contribute to mitigate the impact of anthropogenic warming on coral bleaching. The evidence we present reveals a reduction in bleaching probability at mangrove associated sites when DHW are elevated (Fig. 2). This reduction in bleaching probability is likely to be the result of the significantly higher DHWs threshold which triggers the onset of severe bleaching at mangrove sites compared to the non-mangrove sites (Fig. 3). Therefore, it is likely that corals within mangrove associated reefs have an increased resistance to elevated thermal stress before the most severe levels of bleaching occur. Such increased resistance could result from a multitude of ecological and evolutionary processes, including the preconditioning of corals in the harsher conditions associated with mangrove lagoons (*18*, *23*) and nearshore reef environments (*27*). Furthermore, the different community composition of corals between multi-stressor environments (such as mangrove lagoons) and open water reefs will also effect the resistance of bleaching to thermal stress (*18*, *22*, *23*, *28*), especially given that bleaching responses vary interspecifically (*29*). Or, mangrove associated reefs may be physically buffered from thermal stress compared to open coral reefs (*17*). However, owing to the 4km distance use in this study, preconditioning of mangrove associated coral reefs is more likely than physical buffering.

### Mangroves as potential refuges

The reduction of bleaching occurring on mangrove associated coral reefs, driven by increased resistance to the onset of severe bleaching from thermal stress, highlights mangrove associated reefs as potential refugia for corals to survive through climate change. These refugia are critical as the frequency and intensity of thermal anomalies continues to increase because of anthropogenic heating. Therefore, non-mangrove associated coral reefs will continuingly degrade because of mass levels of severe coral bleaching induced by anthropogenic heating, resulting in mass mortality (*3*, *10*, *30*). Subsequently, changes of the benthic composition in non-mangrove coral reef environments, along with habitat homogenisation, will continue as expected (*2*, *3*). Conversely, mangrove associated coral reefs will be less impacted because of higher resistance to thermal stress which induces severe bleaching (Figs. 2 and 3).

The critical role that mangrove forests perform for enhancing coral reef biodiversity and reef resilience is well established (*31*–*34*). While mangrove forests have already been identified as potential refuges for certain corals (*17*), our findings significantly push forward their importance as refugia from thermally-induced bleaching on a global scale. Consequently, coral bleaching resistance may be enhanced from the protection of mangrove forests, adding to the already multifaceted role mangroves perform for enhancing coral reef ecosystem function (*31*, *32*, *34*). However, even for areas of refugia such as mangrove-associated coral reefs, there will likely be critical thresholds where temperatures continue to increase beyond the suitability of these environments (*35*, *36*). Therefore, while mangrove-associated coral reefs are more resistant to temperature induced bleaching, reduction of greenhouse gasses to limit the extent of climate warming remains key (*1*, *9*, *35*, *37*, *38*).

## Methods

### Bleaching data

Bleaching data were collated from Reef Check (*39*) between 2002-2017, giving 9,456 surveys across 3,686 sites covering 76 countries globally (Fig. 1). Reef Check data are collected using established protocols to a high standard by citizen scientist, utilised in previous macroecological studies (*16*, *19*, *40*). Validity of the Reef Check data set is well documented (*41*). From Reef Check data, coral bleaching (% of population), latitude-longitude coordinates, and the date of each survey were extracted. This allowed for accurate extraction spatially and temporally for associated environmental variables. The global extent and comprehensive time scale used for bleaching severity provides a reliable platform to elucidate environmental variables associated with bleaching severity. The long time series used reduces the likelihood of temporally anonymous bleaching events from causing bias. Furthermore, the global extent also reduces the likelihood of local environmental variables not considered in this study from driving associations between bleaching severity and environmental variables.

### Degree heating weeks data

The global standard to determine thermal anomalies are degree heating weeks (DHW) which quantify the temperature exposure above a 1°C for the local mean climatic temperature at that period in time over the last 12 weeks (*2*, *42*, *43*). The DHWs data were collected to assess the interaction effect between high thermal stress and mangrove area on coral bleaching severity. Weekly DHW values were obtained from the Coral Reef Temperature Anomaly Database (*43*) (CoRTAD Version 6). These data are supplied from the National Oceanic and Atmospheric Administration (*43*) specifically to predict bleaching intensity at a ∼4.6km resolution at the equator. Data were extracted from January 2002 to December 2017.

### Associated mangrove area data

Mangrove data were extracted from the CGMFC-21 MFW global Mangrove database (*44*) version 3 (*45*). The year 2010 was selected being the median year for when coral bleaching data were collected. The CGMFC mangrove database covers mangrove areas globally at a 30m^2^ resolution from the years 2002 – 2012. For full detail of the database creation see Hamilton & Casey (*45*). These data are stored in WGS84 coordinates which can be spatially associated with the Reef Check bleaching data based on geographical coordinates. Mangrove area within a 1, 2, and 4km radius of the surveyed reefs were extracted using the Raster package (*46*) in R Studio 4.0 (*47*).

### Statistical analyses

The effects of mangrove area, DHW and their interaction were modelled using a Bayesian hierarchical ordinal regression (*19*). The hierarchical model allowed for inclusion of spatial variation patterns associated with coral ecoregions (*48*). These ecoregions show internal consistency in their taxonomic configuration, dispersal and isolation processes, and patterns of evolutionary history (*49*). The model is defined by the equation

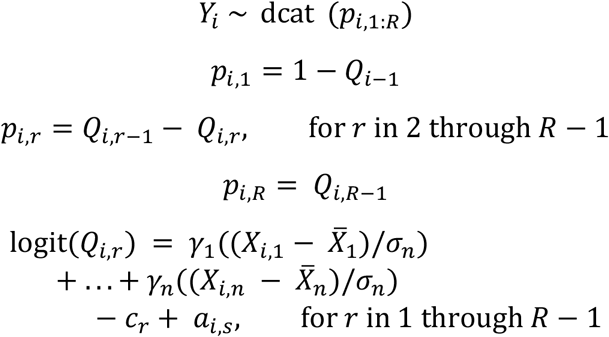

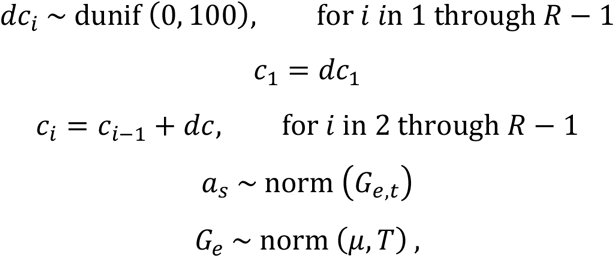

with *Y_i_* being the level of coral bleaching for survey *i* on an ordinal scale (four bleaching severity categories); dcat is a categorical distribution, dunif is a uniform distribution; *R* represents the number of bleaching categories; *p_i_*, 1:4 is the probability of survey *i* being in each of the 4 bleaching categories; *γ* are coefficients, *X* are environmental covariates; σ are standard deviations of environmental covariates; *n* is the number of covariates; *a* are random effects of site (*s*) which hierarchically follow a normal distribution (norm) from the random effect (*G*) of ecoregion (*e*) with mean *μ*; *_t_* is the variance across site; *T* is variance across ecoregion; *Q_i_*, *_r_* is the cumulative probability of having a score of *r* or more; *cr* is a cut point for each category *r*; *dc_r_* is the difference between cut points (*19*, *50*).

The 4 levels of bleaching severity were categorised as (1) no bleaching, (2) mild bleaching (1%-10%), (3) moderate bleaching (11%-50%) and (4) severe bleaching (>50%) (*19*). These data follow a natural ordering and therefore are assumed to represent underlying continuous data (*19*, *50*). In the model, DHW, natural logarithm of mangrove area, and their interaction, were standardized to enhance model stability and incorporated into the model. Flat normal priors were used for the covariates. The Bayesian model was implemented in R studio 4.0.4 (*47*) using the Just Another Gibbs Sampler (JAGS) R interface package “rjags” (*51*) initially with 4000 burn-ins, 5000 iterations, and 3 chains. Convergence of the model was subsequently achieved using the *autojags* function from the “R2jags” package (*52*). Convergence of the model was determined by a Gelman-Rubin statistical value of 1.1 (*53*). Model coefficients were considered to have a significant effect when the 89% credible intervals (CI) did not pass zero (*54*), which is mathematically more stable than using the frequentist 95% threshold when conducting Bayesian analysis with less than 10,000 samples (*55*, *56*).

We ran sensitivity analysis using the Bayesian model described above to determine which distance showed the highest effect. Distance refers to the radius from which the entire area is sampled to determine the total mangrove area around the Reef Check survey. We used a doubling sequence from 1km to 4km (i.e., 1km × 2, 2km × 2) and determined the 4km distance showed the strongest effect for reducing bleaching under thermal stress (Table 1). Differences in bleaching severity thresholds for DHWs were statistically analysed using likelihood ratio tests on the DHW gamma distributions (*16*).

## Acknowledgments

JVJ is funded by the Department for Economy (DfE) Northern Ireland.

## Author contributions

JVJ and DPD designed the study, JVJ collected data and performed data analysis. JVJ, DPD, and JTAD interpreted results and wrote the manuscript.

## Data and materials availability

All Data, codes, and materials used in this study are available on our github at https://github.com/JackVJohnson/Coral_bleaching_mangroves

